# TrAQ: a novel, versatile, semi-automated, two-dimensional motor behavioural tracking software

**DOI:** 10.1101/2024.05.05.592556

**Authors:** Davide Di Censo, Ilaria Rosa, Brigida Ranieri, Tiziana Di Lorenzo, Marcello Alecci, Tiziana M. Florio, Angelo Galante

## Abstract

We present TrAQ, a new MATLAB-based two-dimensional tracking software for Open Field video analysis of unmarked single animal, featuring minimum user intervention. We developed TrAQ with the purpose to automatically count the in-plane rotations, an important parameter in the 6-hydroxydopamine hemiparkinsonian rat model and in many rodent models of neurodegenerative diseases, a very time-consuming manual task for highly trained human operators. In addition, TrAQ allows automatic recognition of the animal within a user defined arena providing a quantitative measurement of the body centroid and the two extremities positions. A full range of quantitative kinematic behavioral parameters are automatically calculated, and the optional shape erosion feature adds usage flexibility. TrAQ, free and non-species-specific application, was quantitively tested with rodents and on a qualitative basis with zebrafish, and invertebrate animal models. Quantitative results were successfully validated against a commercial software (for tracking) and manual annotation (for rotations in an hemiparkinsonian rat model). This is a widely used model in preclinical research to study postural instability and motor asymmetry. TrAQ allows the characterization of motor asymmetry using non-invasive tools, thus appreciating the spontaneous Open Field behaviour of unmarked single animal, with minimum user intervention.

## 1 Introduction

Video tracking analysis of freely behaving animals offers a powerful tool to noninvasively capture a range of quantitative behavioral events, e.g. animal position, speed, posture, activity, social interactions, number of visits in a sub-arena region. Video tracking analysis is much simpler and effective than other tracking methods, such as RFID tags (Lewejohann et al., 2009), GPS (Hebblewhite & Haydon, 2010; Van De Weerd et al., 2001) or accelerometers (Koniar et al., 2016), and does not subject animals to additional stressors such as radio collars or subcutaneous microchip implantation. In addition, video tracking analysis is the only suitable method to track the movement of sub-millimetric animals such as, for example, insect larvae or microcrustaceans (Di Lorenzo et al., 2019). Thus, computer-vision based software for animal tracking is a key complementary toolkit to ethological (Di Lorenzo et al., 2019), genetic, phenotype, molecular and electrophysiological characterizations.

Several commercial video tracking software packages are available, each one with peculiar characteristics and limitations. For the analysis of rodent models, we can list Videotrack (Viewpoint Behaviour Technology, Civrieux, France), EthoVision (Noldus et al., 2001), TopScan (CleverSys Inc., Reston, VA, USA), Smart Video Tracking (Panlab, Harvard Apparatus, Barcelona, Spain) and AnyMaze (Stoelting Co., Wood Dale, IL, USA), all of them considered as gold standards. For aquatic animals standard tracking software suites are Lolitrack (Loligo® Systems, Viborg, Denmark) and the Microimpedance Sensor System (Limco International GmbH, Konstanz, Germany), suitable for aquatic micro invertebrates. These commercial solutions are stable and versatile but can be only used with a restricted number of animal models, may require massive user interaction to setup a study and quite often are sold as hardware and software bundles, adding a significant running cost to behavioral research laboratories.

Several free custom-made tracking software are also available, enabling behavioral measurements of individual (Ben-Shaul, 2017; Crispim Junior et al., 2012; Gomez-Marin et al., 2012) or multiple animals (Giancardo et al., 2013; Pérez-Escudero et al., 2014; Rodriguez et al., 2017; Walter & Couzin 2021; Chen et al., 2023) both in the presence of markers or in the more appealing marker-less modality (for a complete overview see Panadeiro et al., 2021 and Luxem et al. 2023). However, only a few of them were quantitatively validated against commercial tools (Bello-Arroyo et al., 2018). In addition, free software tends to be heavily application-oriented and, unfortunately, quite often the quantitation features of locomotion and exploration behaviors are limited. They may restrict the measure to gait (Leroy et al., 2009; Liang et al., 2012; Lecomte et al., 2021), position (Koniar et al., 2016) or specific postures (Farah et al., 2013; Ou-Yang et al., 2011). Moreover, they can work only with specific laboratory arenas (Aguiar et al., 2007; Korz, 2006) or home cage set-ups (Heredia-López et al., 2013; Koniar et al., 2016). The most recent and advanced use Artificial Intelligence approaches to study animal pose and social behavioral features (Gerós et al., 2020; Mathis et al., 2018; Sun et al., 2021; Pereira et al., 2022; Bordes et al., 2023), which allow impressive performances, at the expenses of being restricted to the animal model used to train the algorithm and, often, to non-straightforward installation and operation procedures. In spite of such limitations open-source tracking software can be widely used and supported by an active community of users and developers, so that the software packages are adaptable and expandable to future assays by including novel behavioral parameters, models, organisms, and experimental set-ups (Nema et al., 2016), although advanced programming skills are necessary, this being a real bottleneck for most biological science labs (Reiser, 2009).

Previously, free behavioral tools have been developed in the MATLAB environment (The MathWorks, Inc., Natick, MA, USA), allowing tracking of single rodents (Ben-Shaul, 2017; Ohayon et al., 2013) or other animals, with some of them being able to track individual animals in a group (Pérez-Escudero et al., 2014). This methodological choice requires a MATLAB license, often already available in academic or research institutions, making the overall investment less expensive than purchasing specialized commercial tracking software. Moreover, the commercial MATLAB user-friendly desktop environment allows an easy software installation and execution. From the end-user side, we notice that MATLAB code is totally platform-independent, compared to other common options like Python or C.

Our interest in video tracking software started with a 6-hydroxidopamine (6-OHDA) hemiparkinsonian rat model (Ungerstedt, 1968), one of the most useful tools in Parkinson’s Disease (PD) research given its high reproducibility and, good characterization, and the possibility to use the intact hemisphere as an internal control (Florio et al., 2018). Unilateral lesions of the nigro-striatal dopamine (DA) system induce a profound asymmetry in motor performance, expressed in rodents as asymmetric body posture, impaired use of the contralateral forelimb, and a sensori-motor orientation deficit (Björklund & Dunnett, 2019). The rotational test is the most widely used and best characterized in hemiparkinsonian rodents because of its simplicity and sensitivity (Ungerstedt & Arbuthnott, 1970). Rotations in 6-OHDA lesioned rats can be easily evoked by administration of DA agonists and are quantitatively related to the degree of DA receptor stimulation (Ungerstedt & Arbuthnott, 1970). The evoked rotational behavior can be automatically registered as the number of complete 360° turns per minute in a commercial rotameter device (*e*.*g*. TSE-Systems or PANLAB), which is considered the gold standard to measure circling behavior of animals. This testing paradigm is effective in rotations counting but does not allow, in the same testing session, to evaluate other locomotor and behavioral observables. The rotation counting in an open field arena is, for the tested animal model, relevant because corresponds to highly demanding experimental conditions, since the apomorphine-induced condition is characterized by a compulsive rotational behavior that heavily affects the usual 2D animal shape.

In such a scenario, we acknowledged the need for a low-cost open-field tracking tool, for users with basic IT skills, allowing automatic in-plane rotations counting for PD and other neurodegeneration studies (Ungerstedt, 1968; Björklund & Dunnett, 2019; Prasad & Hung, 2020), as well as to quantify other behavioral parameters during the same test session. To this purpose we developed TrAQ: a novel MATLAB-based, versatile and semi-automated tracking software able to analyze videos of single-animal motor behavior with minimal user interaction. As better described in the Supplementary Material, the implemented algorithms are based on video intensity thresholding, shape erosion, cluster extraction, animal centroid and body extremities identification. The software allows automatic detection of an unmarked individual animal within an arbitrarily shaped arena, providing 2D coordinates of the animal centroid and extremities (e.g., nose and tail for rodents), as well as kinematic variables useful to quantify motor performance: resting/activity rate, exploration/exploitation activity adding the number of in-plane rotations, to better characterize animal’s circling behavior. Software validation was performed with a set of smartphone-recorded videos of hemiparkinsonian rats, comparing the results with a commercial software (EthoVision XT 130.1220), as well as with manual rotation counts. More qualitative tests of general TrAQ features were performed with, mice, zebrafish and invertebrate copepods.

## 2 Methods

### 3 6-OHDA rat model

We quantitatively tested TrAQ with male albino Sprague-Dawley rats unilaterally injected into the right Substantia nigra pars compacta with 6-OHDA (8mg/4ml) as a hemiparkinsonian model (Ungerstedt, 1968; Casarrubea et al., 2019), following the same surgical procedure described in Rosa et al., 2020. The experimental protocol for animal care and handling was according to the European Community Council Directive (86/609/EEC) and the Italian law n. 26 (14/03/2014) and was approved by the Institutional Review Board of the University of L’Aquila and approved by the Italian Ministry of Health (authorization number 934/2017-PR). 6-OHDA induced unilateral degeneration produces an asymmetric and quantifiable motor behavior in the presence of systemic administration of apomorphine, a dopaminergic receptor agonist. Apomorphine stimulates supersensitized receptors in the lesioned striatum and the induced rotations (up to 800 in a 60 min test) are proportional to the degree of neurodegeneration degree (Ungerstedt, 1968; Blandini & Armentero 2012).

### 4 TrAQ software

TrAQ was developed in MATLAB to avoid incompatibilities among different Operative Systems allowing easy data processing (outputs are also available in Microsoft Excel format). Besides the basic MATLAB installation, the tracking software requires the MATLAB Image Processing toolbox. TrAQ is composed by two main software modules, named Tracker and Analyzer, and their Graphical User Interfaces (GUIs) that allow the operator to control the workflow and data output.

#### Tracker Module

The Tracker includes the following routines: background extraction for frame-by-frame subtraction(optional), pixel intensity thresholding to generate a binary image, binary image erosion (optional), clusters identification within the binary image, largest cluster centroid and extremities identification by geodesic distance (Soille, 2004). Technical details can be found in the Supplementary Material.

After thresholding, a pixel cluster analysis on the binarized image is performed (Fig 1a, 1b), and the largest cluster is associated with the animal. From the largest detected cluster, the Tracker extracts the animal’s 2D centroid coordinates (Fig 1b), the first extremity point (the cluster point with larger geodesic distance from the centroid, the tail-end for rats, Fig 1c) and the second extremity point (the cluster point with larger geodesic distance from the first extremity, the head for rats, Fig 1d). This approach assumes that the animal is the largest moving object in the arena.

**Fig 1.**
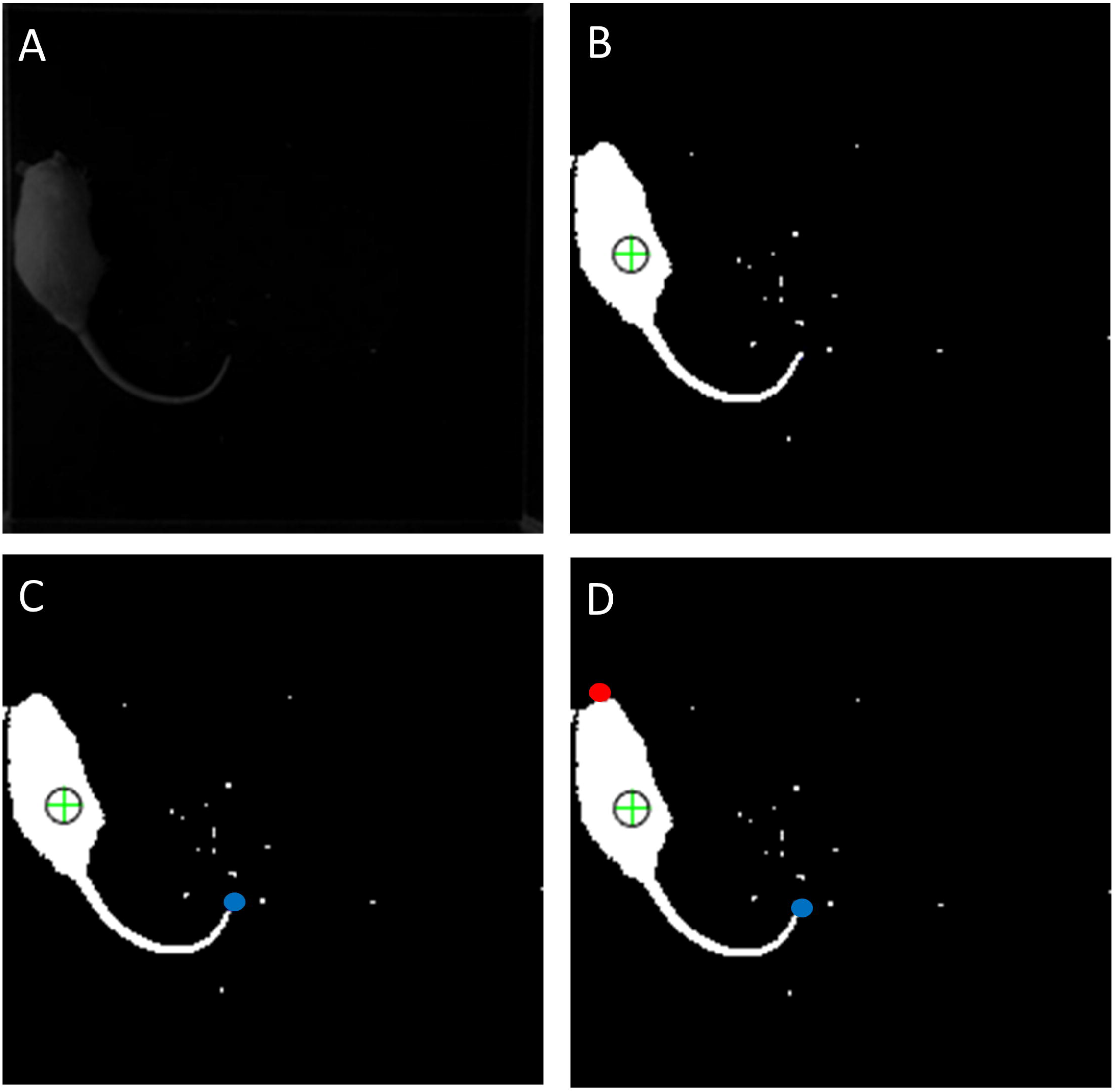
The frame-processing pipeline: **(A)** original image frame; **(B)** binary image after thresholding and largest cluster centroid identification (green cross); **(C)** tail-end (blue point) identification as geodesic farthest point from centroid; **(D)** head (red point) identification as geodesic farthest point from tail-end.

While the centroid position is a very robust datum, the head/tail extremities detection algorithm is sensitive to the shape of the main cluster, i.e. the animal’s posture. We did not detect any problem in extremities localization with copepod and zebrafish videos since these organisms present limited changes in their body posture over time. In rodent models however, posture can alter the mutual distances between the centroid and body extremity points, giving rise to wrong identification of the latter. This is evident during the compulsory behavior of 6-OHDA lesioned rats after apomorphine treatment. The rat tail becomes a confounding element for extremities localization and rotation counting. These localization errors were generated by the recurrent condition of partially visible tail, as in Fig 2b, because positioned under the body during compulsory pivoting and/or grooming. In this condition tail removal resulted beneficial, allowing to restrict the analysis to the main body only, thus extracting its correct spatial orientation (Fig. 2d). Tail removal can be achieved by applying, before the clusters’ identification, the erosion procedure on the binarized image. The erosion eliminates a user-selected pixel thickness from the boundary of each region with above threshold pixels (van den Boomgaard & van Balen, 1992), and can be adjusted to remove animal-specific features (tail, legs) with size smaller than the main body. Cluster erosion can also speed up the video analysis. Indeed, clusters identification is the most time-consuming processing step of the Tracker module. If computing resources are limited, when analyzing large video recordings, a balance needs to be found between speed (high intensity threshold, few clusters, possible loss of part of the animal shape) and precision (low intensity threshold, more clusters, more accurate animal shape and extremities localization). Avoiding long computation times while keeping a satisfactory localization accuracy can be achieved by applying the erosion procedure to eliminate the small non-relevant clusters (typically the ones related to camera noise, lighting conditions changes, animal’s feces or litter residues generated during the test). An illustrative example of clusters number reduction and x10 TrAQ speed increase for a Copepod video is shown in Figs S2a, S2b of Supplementary Material file.

**Fig 2.**
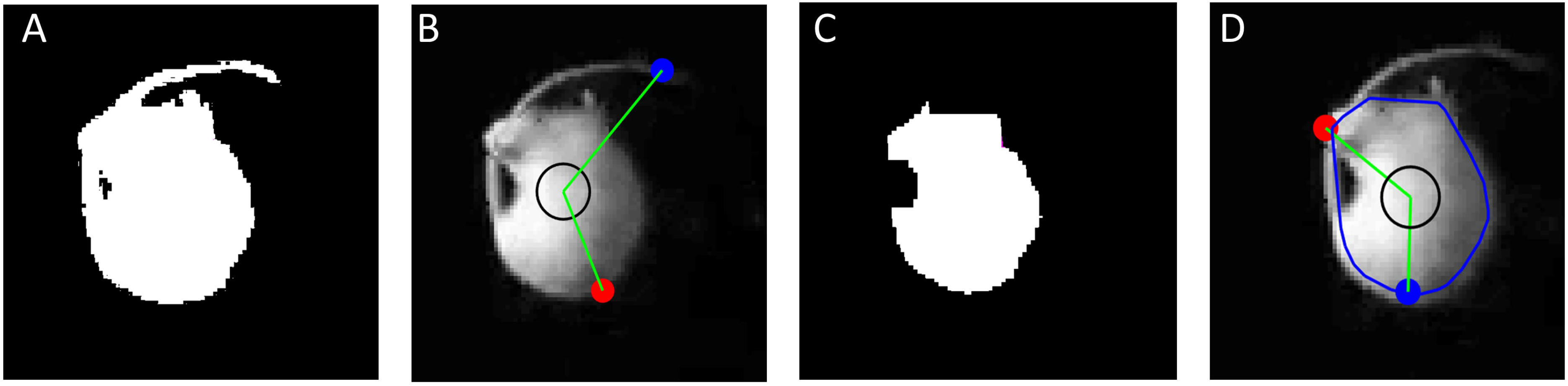
Erosion: details of a PD rat model during apomorphine-induced rotation with altered posture: **(A)** binary non-eroded image without erosion. **(B)** Without erosion extremities localization (colour code as in Fig 1) can be faulty. **(C)** Binary image after applying a 4-pixel erosion procedure, the analysis can be restricted to the main body only (the blue line is the main body convex hull after erosion), with improvement in extremities localization accuracy **(D)**.

The Tracker can be applied to the entire Field of View (FOV) of the video or to a limited portion defined by the user (the animal arena). The latter possibility is particularly useful if (i) the experimental setup presents image artefacts, like reflections of the animal on the arena walls, which may give rise to animals’ shape artefacts; and (ii) there are multiple arenas in the video and a dedicated Tracker run can be performed for each of them.

#### Analyzer Module

The Analyzer takes the Tracker output and allows raw data manipulation to generate the final outputs. The Analyzer module converts position from pixel units to physical spatial coordinates and calculates all derived observables from the tracked positions of centroid, head, and tail. This module also calculates the main body spatial orientation θ, defined as the angle between the video horizontal axis and the oriented segment defined through the animal’s body centroid and the head (Fig. 3). The Analyzer calculates θ for each frame, and its temporal evolution is used to compute the total number of rotations counting. The user can define a centroid velocity threshold to calculate the animal’s activity index, defined as motion-to-rest time fraction. We noticed that, using tail erosion and with largely deformed rat postures, in some frames the extremities detected by the Tracker could be assigned the wrong label, thus the animal head is wrongly classified as tail-base and vice versa. We call this a label swap and their incidence, defined as ratio of frames with swapped extremities divided by the total number of video frames, is a crucial parameter affecting the TrAQ rotation counting accuracy. To solve this problem the Analyzer model can implement two label correction algorithms: one based on a head-on movement assumption, another on the enforcement of the extremities’ spatial continuity criterion between frames. Based on our experience, the spatial continuity criterion offers better performances when the animal does not assume a posture where its head can touch the back of the body (e.g. hind paws grooming or compulsive pivoting in rats), while the head-on movement assumption works better in all the other cases. A full description of these algorithms is provided in the Supplementary Materials file.

**Fig 3.**
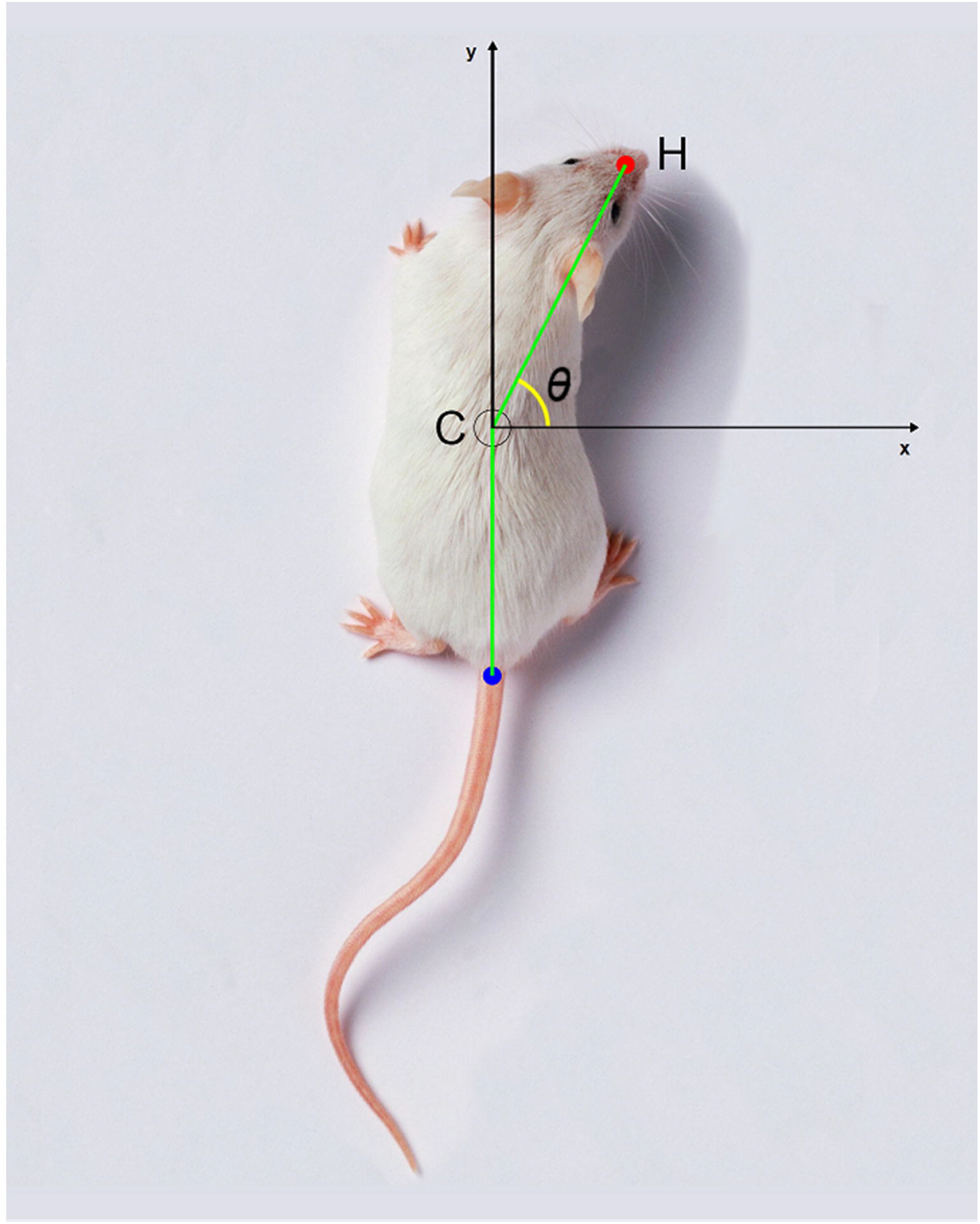
Head direction. The head direction is defined as the angle (*θ*) between the video frame horizontal axis (x) and the segment defined through the animal’s centroid (C) and its head (H).

Any change of the analysis parameters within the Analyzer will generate updated outputs. The whole range of observables is easily accessible in the results file, in MATLAB as well as Excel formats, which includes the frame timestamp. The TrAQ absolute time identification allows the user a posteriori comparison (within the temporal resolution of the video frames) with data recorded by other software packages, tools, and techniques (e.g. electrophysiological data).

Another feature of the Analyzer is the possibility to generate and save the background-subtracted version of the original video. This processed video improves the contrast of the tested animal and can be useful for further visual analysis of skilled operators (see Fig S1a and S1d in the Supplementary Material).

### 5 TrAQ output

TrAQ implements a batch process function: if the user needs to analyze a set of videos taken in similar environmental conditions (e.g. same arena and camera positions, similar lighting levels), it is sufficient to set-up the tracking settings of the first video and the entire batch of videos can be processed without any other operator’s interaction.

Typical examples of output plots from a 10-minute video of a rat recording are shown in Fig 4: the Analyzer’s Results Viewer window allows to select several output graphical formats, such as the centroid and/or extremities track (Fig. 4a), position heatmaps (Fig. 4b), the instantaneous speed plot (Fig. 4c), the corresponding speed histogram (Fig. 4d),, the centroid-head (line of sight) angular orientation histogram (Fig. 4e). The numerical data can be exported as MATLAB (.mat) or Microsoft Excel (.xslx) files for further analysis and graphical presentation.

**Fig 4.**
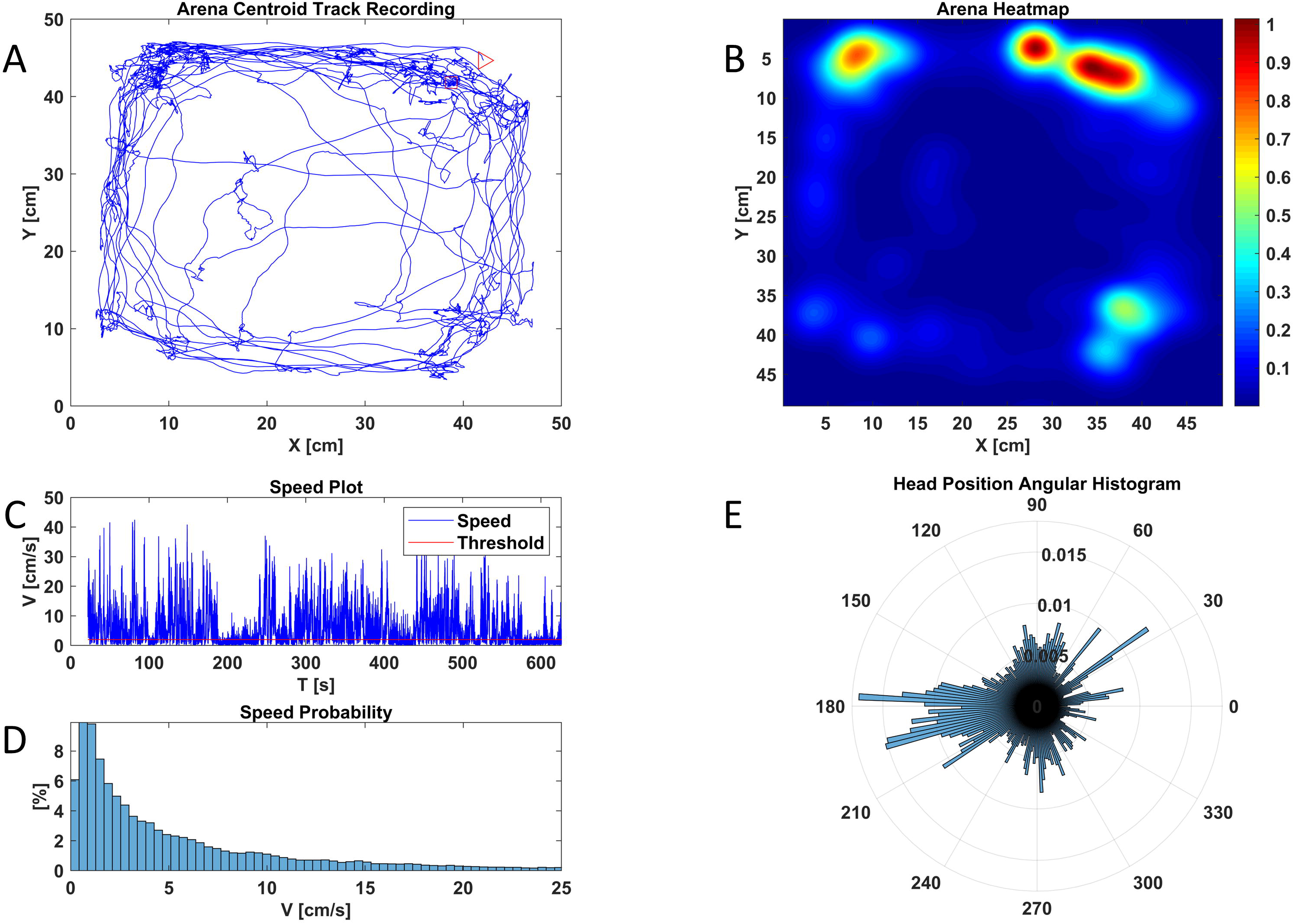
TrAQ graphical outputs. TrAQ graphical outputs of a single rat recording (10 minutes): **(A)** centroid 2D tracking coordinates within the user defined arena (50 cm x 50 cm); **(B)** the corresponding centroid heatmap representation; **(C)** time course of the centroid instantaneous speed; **(D)** corresponding speed probability distribution histogram; **(E)** centroid-head angular orientation (with respect to the horizontal axis) histogram.

The requested CPU time is dominated by the Tracker pipeline, while the Analyzer runs almost in real-time. The computational cost depends on the video resolution and the number of detected clusters. Using a relatively underpowered desktop PC equipped with 4 GB of RAM and i7-4770k processor the typical total computation time for a low resolution (208 x 360, 30 fps) 10 min rat video is 190 seconds, while for a 1 min high resolution (1920 x 1080, 30 fps) copepod video it takes 32 minutes, which are reduced to 3 minutes when the erosion procedure is included (without any appreciable difference in the results quality).

### 6 Validation and Data Analysis with Animal Models

#### Quantitative 6-OHDA rat model

TrAQ was validated against the commercial software EthoVision XT 13.0.122. For quantitative comparison we used the Bland-Altman plots (Bland & Altman, 1986) which allow a compact visual representation of differences in large data sets. We analyzed the videos and compared outputs from twenty videoclips (one-minute long each) from 6-OHDA rats, 15 in drug free condition and 5 after apomorphine administration (Fig. 5). The comparison involved the centroid 2D (Fig. 5a) and 1 D coordinates (Fig. 5b, 5c), as well the total travelled distance (Fig. 5d).

**Fig 5.**
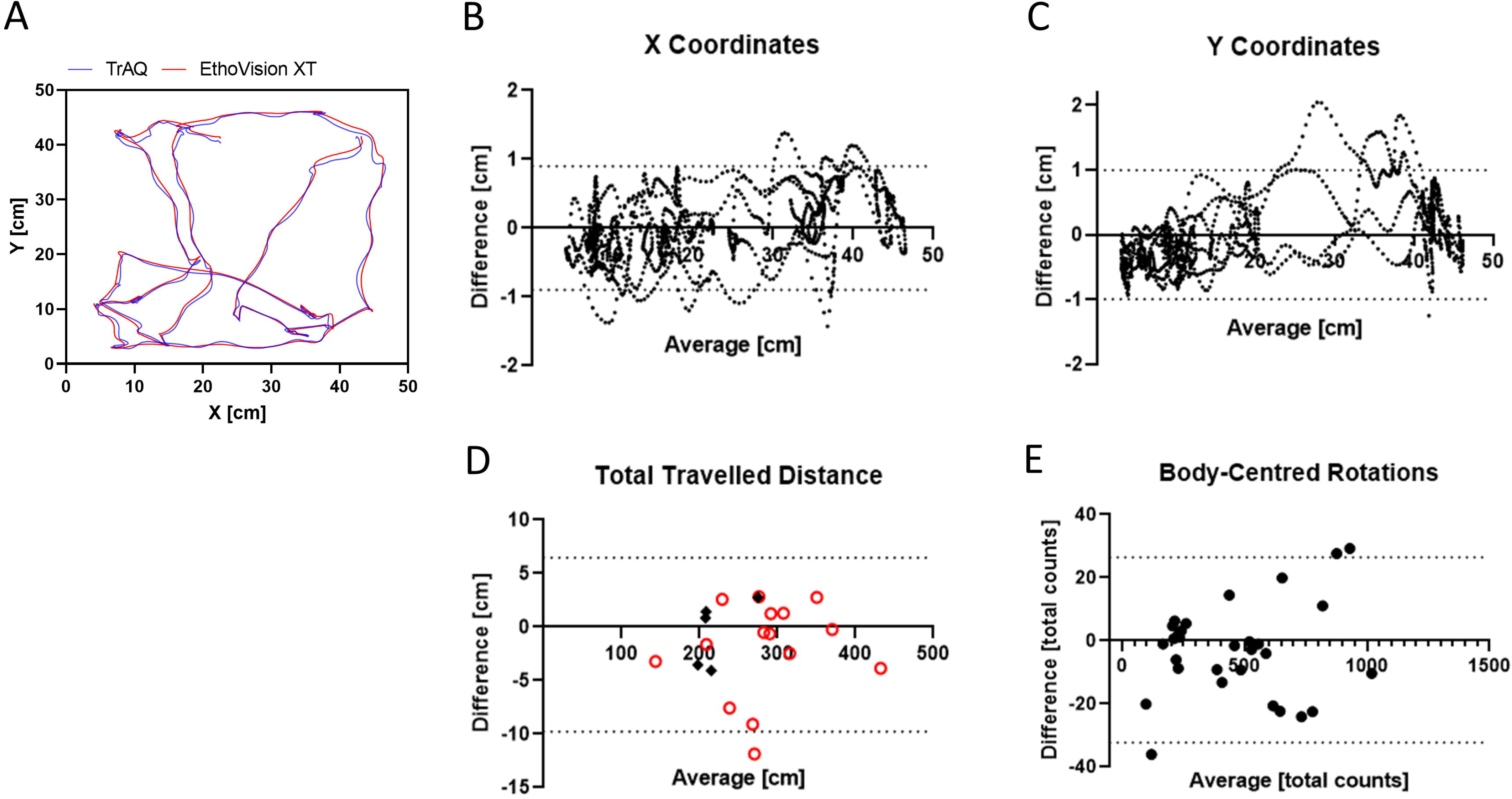
TrAQ results validation. **(A)** TrAQ and EthoVision XT centroid 2D tracking within the user defined arena (50 cm x 50 cm). **(B)** centroid x-coordinate and **(C)** centroid y-coordinate Bland-Altman plots, comparing the same data sets of **(A)** from a one-minute videoclip of a 6-OHDA lesioned rat in drug-free condition; the dashed lines correspond to 95% confidence intervals. **(D)** Total travelled distance calculated from twenty videoclips (one-minute long each) in the rat drug-free (open red circles) and apomorphine-induced (black filled diamonds) conditions. **(E)** TrAQ and manual annotation of the net number of rotations measured in incremental clips ranging from 1 to 60 min in the rat apomorphine-induced condition.

The number of rats’ net body-centered turns, in apomorphine condition, were compared with the manual annotation since neither Ethovision nor other free software provides such feature. To test the rotation counting stability over time (i.e. within a wide range of rotation counts) we used 8 additional videos of different length (1, 3, 5, 10, 20, 30, 40 and 60 minutes) in apomorphine-induced conditions (Fig. 5d).

Of course, a larger number of qualitative and quantitative examples with other species and/or other tracking software tools would be desirable, however we believe that the selected 6-OHDA rat model is one of the most complex and demanding due to the large variations of animals’ 2D shape within the experimental observation window (60 min).

We analyzed behavioral videos of 6-OHDA and SHAM rats divided into five experimental groups: pre-lesion animals (PRE, n=59), Sham and 6-OHDA rats in free (F) condition (SHAM F, n=66; 6-OHDA F, n=108), Sham and 6-OHDA rats under apomorphine (A) challenge (SHAM A, n=56; 6-OHDA A, n=96) recorded with a smartphone at 1920×1080 pixels resolution, using an RGB, 30 fps, .mov file format. The native video resolution corresponds to sub-millimetric pixel size, a spatial resolution usually not needed for such rat model but with a considerable impact on the analysis computational time. Thus, the videos were converted to 360×280 pixels, RGB, 30 fps, .avi format before analysis. The animals were tested several times in a 21-day period after the surgery and the total number of videos was 385 for a total of 287 recorded hours. An example of 2D tracking outputs of sham and 6-OHDA rats is reported in Fig. 6. Statistical comparison between groups was done using the Kruskal-Wallis test with Dunn’s post-hoc.

**Fig 6.**
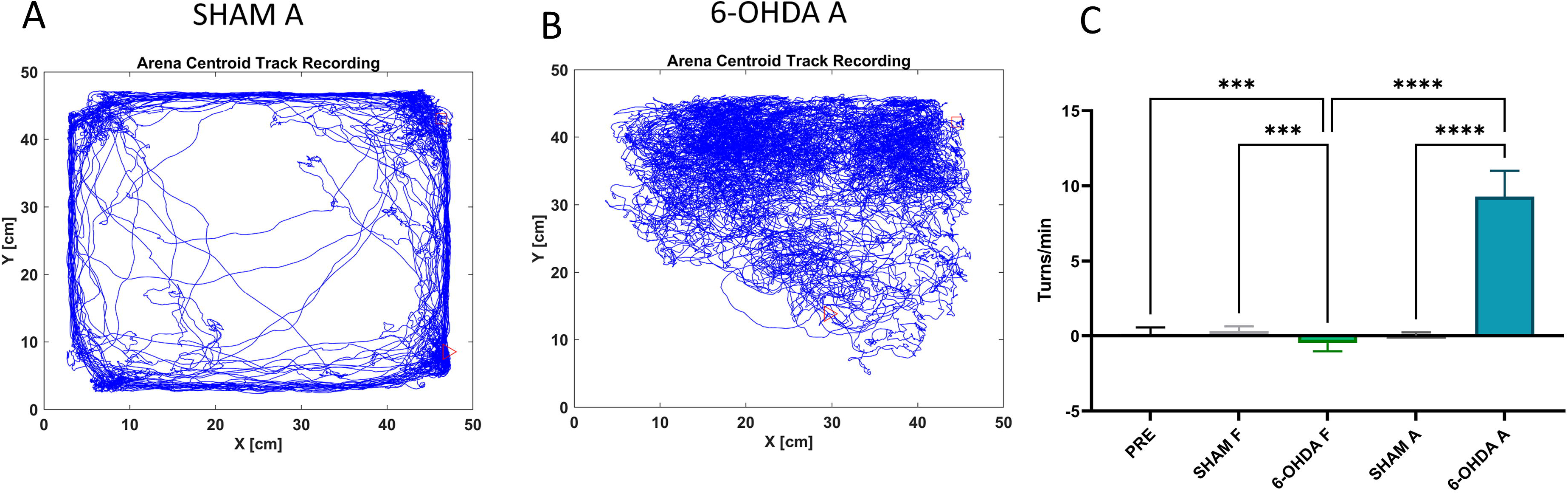
6-OHDA effects. 2D tracking examples of: **(A)** SHAM rat; and **(B)** 6-OHDA rat after APO administration. **(C)** Net number of body centred turns pooled data (ΔR=clockwise-counterclockwise rotations). Intergroup comparison for Pre-surgery (t = 0, PRE) and pooled data of SHAM and 6-OHDA rats in drug-free (F) and apo-induced (A) conditions. PRE vs 6-OHDA F (p<0.001); SHAM F vs 6-OHDA F (p<0.001); 6-OHDA F vs 6-OHDA A (p<0.0001); SHAM A vs 6-OHDA A (p<0.0001). Kruskal-Wallis test with Dunn’s post-hoc.

#### Qualitative mice, zebra fish and copepod model

A previous TrAQ version was used to analyze *Diacyclops belgicus* videos (Crustacea Copepoda) and results were reported in Di Lorenzo et al. (2019).

To illustrate the versatility of TrAQ with respect to species, animal size and spatial resolution, we performed qualitative tests with mice, zebra fish and copepod models. The tracking accuracy was visually verified on the videos, not against other software. We analyzed videos based on the mice Novel Object Recognition paradigm, see Fig. 7b. The experiments were carried with 5xFAD Alzheimer’s disease mouse model at King’s College London under the UK Animal (Scientific Procedures) Act 1986 Amendment Regulations 2012 on Project Licences P023CC39A. We also performed qualitative tests with the zebrafish model, common for laboratory studies on aquatic animals, using internet available videos (Esancy et al., 2018), see Fig. 7c. Finally, TrAQ was used to analyze *Diacyclops belgicus* videos (Crustacea Copepoda) based on previous work by Di Lorenzo et al. (2019).

**Fig 7.**
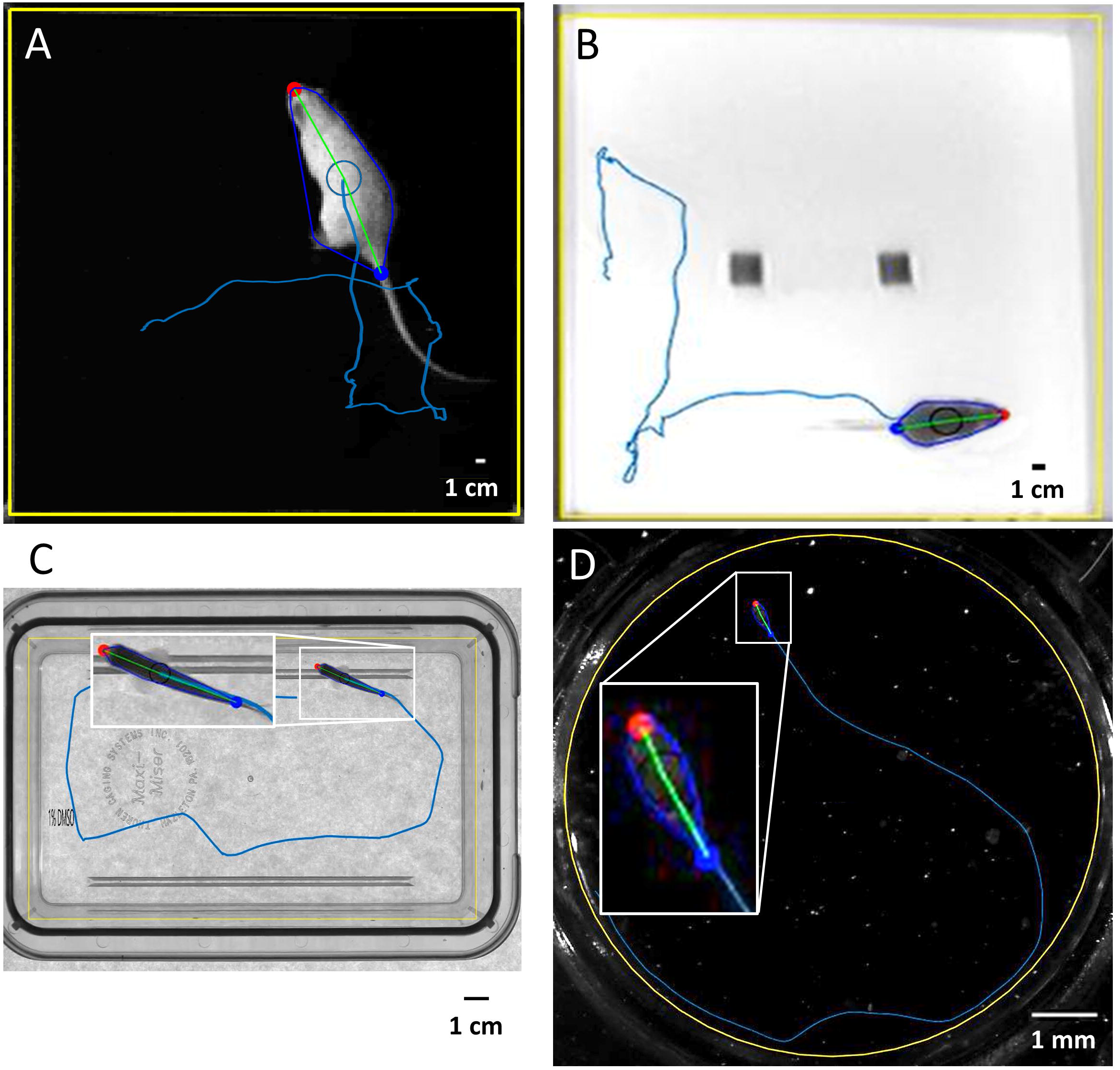
Tracking with different species. Examples of animals tracked in the open field with TrAQ: **(A)** Rat in a square arena of 50 cm size; **(B)** Mouse in a square arena of 40 cm size; **(C)** Zebrafish in a rectangular arena of 17.8 cm x 10.16 cm size. **(D)** An adult individual of the copepod specie Diacyclops belgicus in an 8-mm-diameter microwell observed with a stereomicroscope. The inserts in **(C)** and **(D)** were used to appreciate the successfully track of the head/tail extremities for the small size animal species (zebra fish about 2 cm; copepod 1 mm).

## 7 Results

### 8 Centroid position against EthoVision XT

Both TrAQ and EthoVision XT adopt the centroid’s coordinates to calculate a large variety of secondary behavior parameters (e.g. total travelled distance, speed, activity, spatial occupation frequency). A fair comparison requires an excellent agreement in the coordinates centroid’s derived by the two software, to ensure a high accuracy for the dependent secondary parameters (total distance; spatial probability distribution; velocity; activity; acceleration; probability distribution parameters).

An example of trajectories, calculated on the same video, by TrAQ and EthoVision XT is shown in Fig 4a. In Figs 4b, 4c Bland-Altman plots are shown, comparing tracking results from EthoVision XT and TrAQ for both the centroid x and y coordinates within a time window of 1 minute. We found that the maximum difference of the centroid coordinates between the two methods was typically < 2 % of the total arena linear size (50 cm). We also observed agreement in the total travelled distance calculated from 20 one-minute long videoclips in drug-free and apomorphine-induced conditions (Fig 4d).

### 9 TrAQ automatic rotations against manual counting

We tested TrAQ automatic rotation counting, for 6-OHDA hemi-parkinsonian rats, by comparison with the golden standard manual results obtained by expert operators. The incidence of labels swap (defined as ratio of frames with swapped extremities divided by the total number of frames) is a crucial parameter affecting the TrAQ rotation counting accuracy. We found that, with lesioned rats in drug-free condition (see Fig. 5a for a typical track path) and complete tail erosion, the TrAQ raw data output presented an average of less than 0.5 % extremity’s labels swap. As expected, the same observable was much more challenging in the apomorphine-induced condition (see Fig. 6b for a typical track path), when the animal is compulsory rotating with a high deformation of its top-view shape. In this case the swap rate increased to about 5 %. After the application of the automatic label correction algorithm the swaps were reduced to 0.005 % and 0.04 % respectively, providing a very accurate rotation count against the human operator counting (Fig 5e). The maximum deviation between the two data set was < 3 % of the total number of rotations for the worst-case videos (apomorphine-induced condition). Using the TrAQ rotation counting feature, with an initial effort to fix some parameters (intensity threshold, erosion size), we automated the analysis of the entire study which took about 30 CPU hours on the above-described PC. The results are summarized in Fig. 6c. They provide statistical evidence of the successfully induced damage when the animals are tested in apomorphine challenge (SHAM A vs 6-OHDA A). At the same time, they also show the development of a spontaneous motor asymmetry (SHAM F vs 6-OHDA F and PRE vs 6-OHDA F) which, despite being much smaller than in the previous case, still holds a solid statistical significance. This is the first time, to the best of our knowledge, that such small but statistically significative differences are observed with the 6-OHDA rat model using an automated open-field rotation counting software.

## 10 Discussion

TrAQ was developed having in mind the needs of small-budget behavior research groups finding difficulties in purchasing and maintaining the costs of a commercial product and/or finding not appropriate or difficult to use, for the specific model/test, other available free software. The TrAQ rationale, based on cluster analysis of thresholded frames, makes the package of very general use and suitable for both aquatic and terrestrial species of any size and shape. The built-in erosion algorithm adds flexibility, helping the removal of confounding features from the animal shape (i.e. the tail for rodents) or removing small spurious clusters to reduce CPU time. This widens TrAQ range of applications without the need of advanced programming expertise. The MATLAB-based approach removes software installation complexity, allows a wide compatibility of input video file formats, and guarantees software longevity without maintenance efforts on the user side.

Although the TrAQ experimental validation reported here is based on video recording of rectangular Open Field test arena, the software allows the user to select any arbitrary arena shape (Fig 7). We showed that TrAQ has similar tracking performances, typically within 2 % for the rat centroid coordinates (i.e. less than 10 mm), with respect to the commercial gold standard software (Fig 5a). This centroid tracking data validation gives us a very good confidence in the derived observables commonly used in most behavioral studies.

TrAQ, as compared to the currently available free software packages, adds a few innovative features like accurate rotations counting (Fig 5e), probabilistic background subtraction and easy output data access. To the best of our knowledge, this is the first time that body-centered rotations, considered the gold standard for the assessment of neurodegeneration in hemi-parkinsonian rat models (Ungerstedt & Arbuthnott, 1970), are measured in a fully automated way in an Open Field environment. These results suggest that TrAQ can be used to reduce intra and/or inter-operator errors, providing robust behavioral data. Based on our validation, we can conclude that TrAQ is a useful tool in elucidating between different rotational performances of PD models (Christensen et al., 2018), or between misleading drug-induced effects (Björklund & Dunnett, 2019). Based on the 2D analysis, TrAQ output can be used to evaluate indices of the motor performances such as: clockwise/counterclockwise circling and sensorimotor in-plane orientation defects.

Multiple Region(s) Of Interest (ROIs) within the analyzed arena can be defined *a posteriori* to evaluate the behavior response to specific stimuli. Once the ROIs are defined by the user, the Analyzer automatically calculates the number and duration of visits for each ROI, a feature that gives potential for standard novel object recognition tests as well as for studies of the animal’s exploration behavior (Belzung, 1999; Eilam & Golani, 1989).

The lack of TrAQ specificity, as usual for one-size-fits-all approaches, has its limits. The first one is accuracy. If centroid coordinates are a robust datum, more complex algorithmic approaches can improve the spatial localization of specific shape features (like extremities in rodents) at the cost of a model-specific (and sometimes task-specific) code. Besides centroid localization, TrAQ is currently limited to extremity points detection. Since the extremities’ assignment is based only on geodesic distance criteria, the software occasionally can incorrectly assign the extremities position when the animal shape is highly deformed. Extremities position continuity criteria, with an operator selected maximum allowed displacement within consecutive frames, can be enforced to reduce such errors and provide reasonably accurate rotation counting in rodent models. The second limit of TrAQ is execution time which prevents real-time analysis of high-resolution videos. This could be improved using other advanced programming languages and software tools. However, based on our experience, we found that the execution time is not a practical issue (it takes 2 minutes for a standard desktop PC to analyze a 10 min behavioral video with a task-optimized videos resolution) while the implemented batch capability is a useful tool in speeding up the research activity with a minimum operator intervention.

## 11 Conclusions

We aimed to develop a simple and user-friendly software able to analyze many videos with a single click, thus reducing the time needed to set-up projects while still providing a large amount of quantitative data. TrAQ is a novel, versatile and automated MATLAB tracking software developed to perform basic analysis in a simple way and adaptable to a wide range of 2D behavioral assessment of laboratory animals, both terrestrial and aquatic. It is not based on model-specific features recognition, rather on image contrast and cluster analysis. Any arena shape and size can be used and the user-defined manipulation of the animal shape, via the erosion procedure, helps in adapting TrAQ to different tasks.

TrAQ simple and robust approach turned out successful for different behavior animal models, despite its development was mainly performed having in mind rodent applications. Rodents tracking is challenging, due to the large variety of different postures they can adopt and the huge alterations that can be induced by lesions and/or drugs (Casarrubea et al., 2019). We developed a specific and unique TrAQ functionality that automatically counts the net number of rotations, an important parameter used in many preclinical rodent models of neurodegenerative diseases, not provided by any other software package.

To make the software of simple use, we provided intuitive GUIs and an instruction manual that explains the kick-off steps. TrAQ was developed in a machine-independent programming language that should simplify its use by the wide community of behavioral research groups, at the cost of requiring a software license from an external vendor (MATLAB) often already available in many research institutions.

The current main limitations are: (i) tracking of a single animal in the arena; and (ii) extraction of 2D locomotion variables only. Currently, TrAQ cannot analyze multiple arenas in the same video, but this limitation can be overcome following a multiple step approach, i.e. with multiple runs selecting each time a different arena inside the video FOV.

## 13 Author Contributions

A. Galante and D. Di Censo designed and developed the software; T. Florio and M. Alecci designed the hemiparkinsonian rats study; B. Ranieri, I. Rosa and D. Di Censo performed hemiparkinsonian rats behavioral experiments; I. Rosa and D. Di Censo participated to the mice animal model development; T. Di Lorenzo designed and performed the copepod ecotoxicological study. All authors discussed the results and contributed to the final manuscript.

## 14 Funding

The research was supported by funds provided by the University of L’Aquila (RIA 2016-2018) (A. Galante, M. Alecci, T.M. Florio).

This work has been funded by the European Union - NextGenerationEU under the Italian Ministry of University and Research (MUR) NationalInnovation Ecosystem grant ECS00000041 - VITALITY - CUP D73C22000840006

15 Acknowledgments

The authors wish to acknowledge Dr. Mattia Di Cicco (University of L’Aquila) for helping in analyzing copepod videos, Dr Diana Cash and Dr. Mattia Baraldo (Kings College London, UK) for helping in analyzing mice videos. We also would like to thank Dr. Esancy and colleagues for letting us use videos from their study.

## 16 Data Availability State

TrAQ is freely provided for research purposes only. We encourage behavioral scientists to use and modify TrAQ for their research needs, with the kind request to cite the software and the present article. TrAQ is freely available and downloadable from the Figshare repository (doi.org/10.6084/m9.figshare.9863378.v1). The repository also contains a manual and an example video. TrAQ has been tested with MATLAB releases from 2018a up to 2022b.

