## Supplementary data for "TrAQ: a novel, versatile, semi-automated, two-dimensional motor behavioural tracking software"

SUPPLEMENTARY MATERIAL

*Tracker Module*

In the process pipeline the Tracker converts RGB videos to a standard BW format or the user can select a specific color channel (the latter feature can be useful to maximize the contrast with respect to the background). Currently, the software can analyze mp4, avi, mpg or wmv videos and, if needed, compatibility can be easily extended editing the video_files.m file located in the software main directory.

Unlike other commercial and/or open-source tracking software TrAQ adopts a probabilistic approach to calculate the reference background image by assigning, to each pixel, the most probable value calculated from a random subset of video frames (Figs S1a, S1b). Based on our experience, this approach outperforms the average pixel intensity approach (Fig S1c), often implemented by other software, particularly when the animal shows a static behavior during a significant (but not prevailing) time interval within the video. Background subtraction is not a compulsory step for the subsequent behavior analysis. However, it helps in maximizing animal visibility (from Fig S1a to Fig S1d), usually very low for experiments on nocturnal animal, such as rodents and subterranean animals – whether aquatic or terrestrial - that are better performed in low light conditions to ensure unbiased responses.

**Fig S1. Background subtraction**: (a) original image frame taken from above the arena; (b) background reference image, computed from the video containing the moving animal, using the TrAQ probabilistic approach; (c) background reference image computed with the standard pixel intensity-averaging process (used as default algorithm in most software); (d) TrAQ background-subtracted frame obtained using (a) and (b).

d

c

b

a

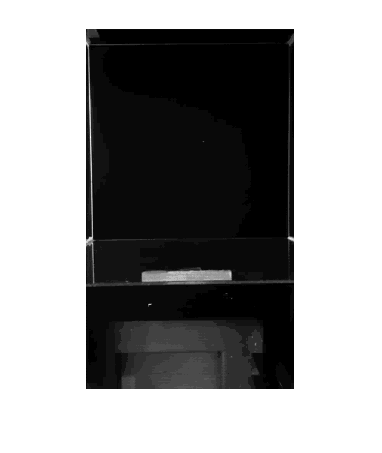

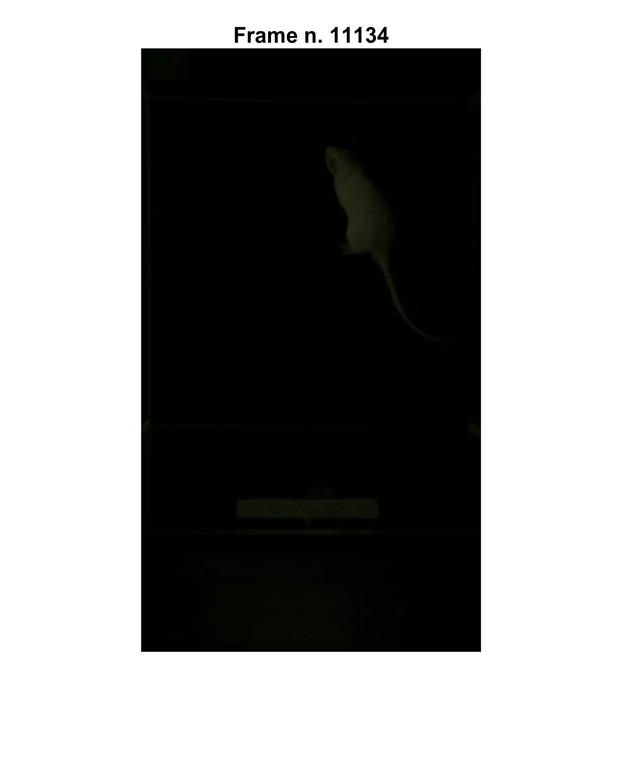

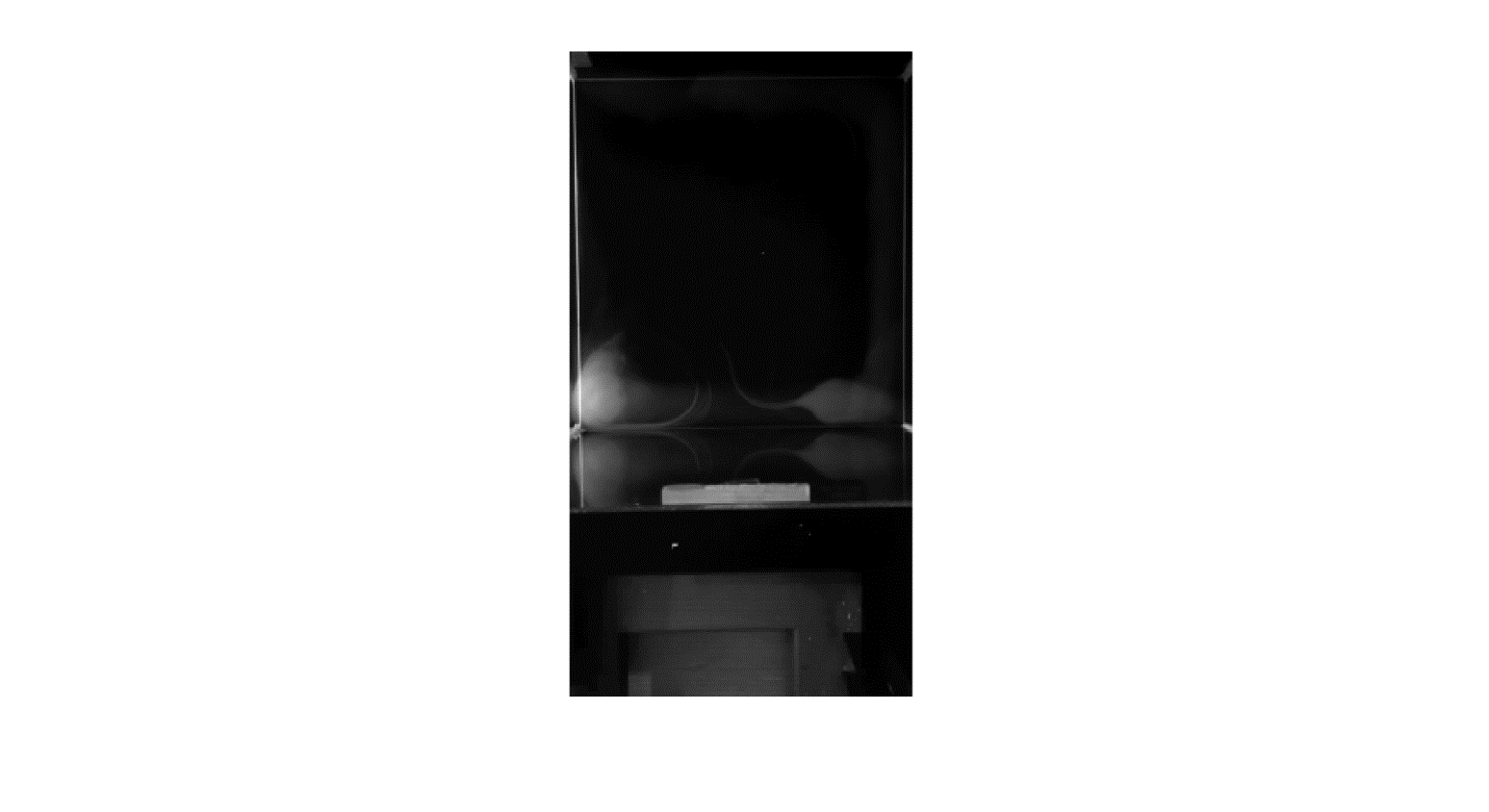

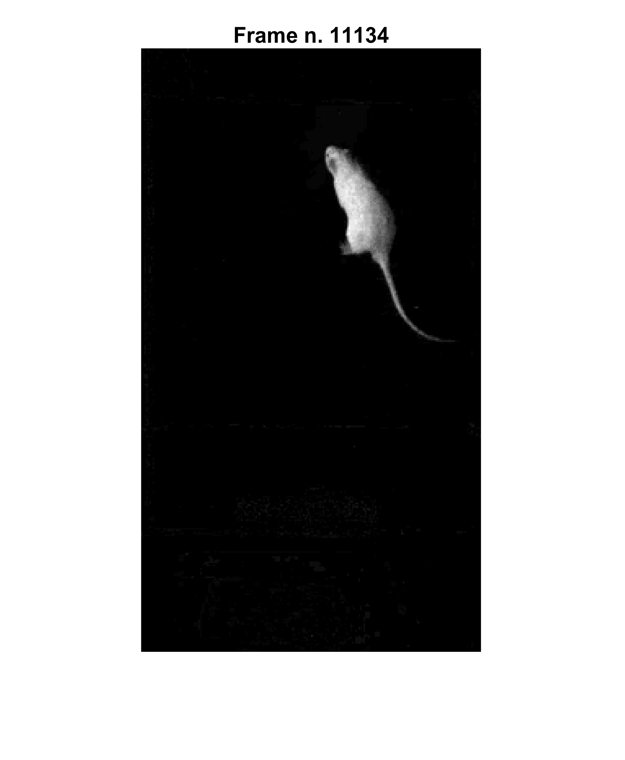

b

d

A pixel cluster analysis on the binarized image is performed, and the largest cluster area is associated with the animal (if needed, in the UserSettings.txt file can be defined an optional threshold on the minimal acceptable size for the largest pixel cluster size). From the largest detected cluster, the Tracker extracts the animal’s 2D centroid coordinates, the whole-body area, and the cluster convex hull. The animal’s extremities coordinates (tail and head) are calculated as specific pixels within the main cluster frontier defined using the geodesic distance: the tail-end is the farthest point from the centroid, the head is the farthest point from the tail-end (Soille, 2004).

The erosion is also a useful tool included into Tracker to increase the analysis speed if a large number of clusters are present in the binarized image. Clusters number is a critical factor for the execution speed of the detection routine. The erosion can be used to wash out the small clusters, thus speeding up the most CPU-intensive part of the Tracker computations. An illustrative example of clusters reduction by adopting the erosion strategy is shown in Figs S2a, S2b which refer to a Copepod video (Di Lorenzo et al., 2019).

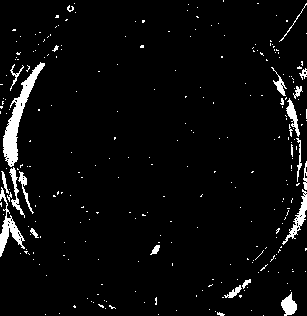

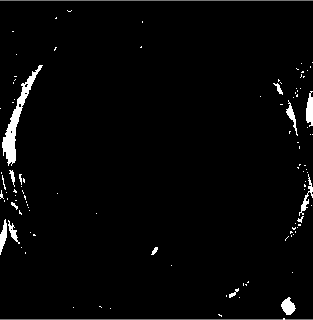

a

b

**Fig S2. Erosion**: binary image (from a video recorded with a stereomicroscope at x12 magnification) of a Copepod (inside the green circle) in an 8-mm-diameter microwell arena (yellow circle) filled with water. (a) The debris have comparable intensity level and show up in the binary image as small clusters. (b) A 3-pixel erosion algorithm removes most debris clusters in the ROI (Region of Interest) and speeds up the analysis by a factor of 10, with some reduction of the effective copepod size.

If, in a frame, the algorithm is unable to track the animal (i.e., to assign the 2D coordinates of centroid, head and tail), the frame is labelled as “untracked”, and the analysis advances to the next frame. When untracked frames are present, the operator can manually fine-tune the intensity and cluster size thresholds to recover them. In case only few frames, with respect to the total number, are untracked this time-consuming option (the Tracker must reanalyze the entire video) can be avoided, and their output may be recovered in the next Analyzer processing step.

*Analyzer Module*

If untracked frames are present in the Tracker output, the Analyzer can fill the gaps by a (linear) interpolation of the animal coordinates (centroid, head, tail) using the closest previous and subsequent successfully tracked frames.

Even with the adoption of the appendix’s erosion algorithm (i.e. to remove the tail in a rodent model, Fig. S3a), extremities identification failure is still possible. Head and tail-base detection failures are due to: (i) misassignment within the set of animal’s cluster border points as in Fig. S3b (it occurs for highly deformed animal postures); or (ii) extremities labels swapping (in Fig. S3c the red point should be associated to the head, not to the tail-base). The latter produces on data an effect analogous to a rapid (within the frame refresh time) $\pm$180^o^ rotation of the animal and, in case of several occurrences, can interfere with an accurate count of rotations.

The Analyzer allows two different approaches for extremities misassignments correction. The first is based on a spatial continuity criterion to re-assigns the head and tail labels to the extremity points identified by the Tracker: the head and tail are respectively the extremity points of the new frame closer to the head and tail in the previous frame. The user also has the freedom to change, in the GUI interface, the head assignment in a frame and the spatial continuity criteria will be enforced in all the subsequent ones. The second approach is totally automatic and works well for the animals that usually proceed only in forward direction, i.e. it requires accordance between the animal’s movement direction and the centroid-head position. The Analyzer calculates the dot product between the vectors that connects the animal’s centroid (C) and head (H) in frame *i* and in frame *i-1*: if the animal has a forward movement $D=\vec{CH}_{i}\cdot\vec{CH}_{i-1}$ should be positive. If D is negative, a head misassignment is assumed and the extremities labelling performed by the Tracker module for frame *n* are reassigned swapping them.

The final user can choose which algorithm to adopt for correction of extremities assignments, based on the animal model under study, directly from the software GUI (Auto Fix and Manual Fix buttons).

To have a large impact on net rotations count the swaps should be comparable with the number of rotations and, in our experience, this never happened. If rotations count is our goal, we suggest using first the Auto Fix procedure. If necessary, a fine tuning of the results can be done, in an interactive session, with the Manual Fix, correcting the few residual errors.

The Analyzer performs rotation counting from the body angular orientation $\theta_{i}\in\left[ -\pi,\pi\right]$ calculated for each video frame (where i = 1, ..., M labels the video frames). The body angular orientation ($\theta_{i}$) is defined as the angle between the video’s horizontal axis and the animal’s centroid-head oriented segment. The time succession of raw angles has 2π radians jumps between consecutive values when the centroid-head orientation crosses the negative *x-*axis orientation. This effect is corrected by an "unwrapping" algorithm which adds multiples of ±2π to $\theta_{j\geq i}$ until $\Delta\theta_{i}=\theta_{i}-\theta_{i-1}\leq\pi$. After the unwrapping procedure, dividing the last frame value $\theta_{M}$ by 2π we obtain the net rotation count (clockwise minus counterclockwise) of the entire video. Even after the unwrapping procedure |$\Delta\theta_{i}|$ can still present jumps smaller that 2π but too large to be compatible with the rotational velocity of the animal. This happens when errors in the extremities localization of a single frame translate in large error in the body angular orientation. Such errors should sum to zero in a video with many frames, when the animal rotates in both directions. But they can provide a non-zero average when there is a preferred rotation direction, as for the hemiparkinsonian 6-OHDA model. To improve the accuracy of the rotation count we introduced a physical constraint on the instantaneous rotational velocity of the animal. If, after unwrapping, |$\Delta\theta_{i}|$ is larger than a user specified threshold $\Theta$, the algorithm checks the animal orientation in a subsequent but temporally close frame with index $n=i+N$calculating $\Delta\theta_{i,N}=\theta_{i+N}-\theta_{i-1}$ with *N* a user defined parameter. If $\Delta\theta_{i}>$ $\Theta$ but $\Delta\theta_{i,N}<\Theta$ , $\theta_{i}$ is probably affected by a large error in extremities localization of frame *i* and the unrealistic jump is cancelled redefining $\theta_{j\geq i}=\theta_{j}-\Delta\theta_{i}$. For 6-OHDA hemiparkinsonian rat model under apomorphine challenge we adopted $\Theta=4/3 \pi$ and N=10 (corresponding to 0.3 s).

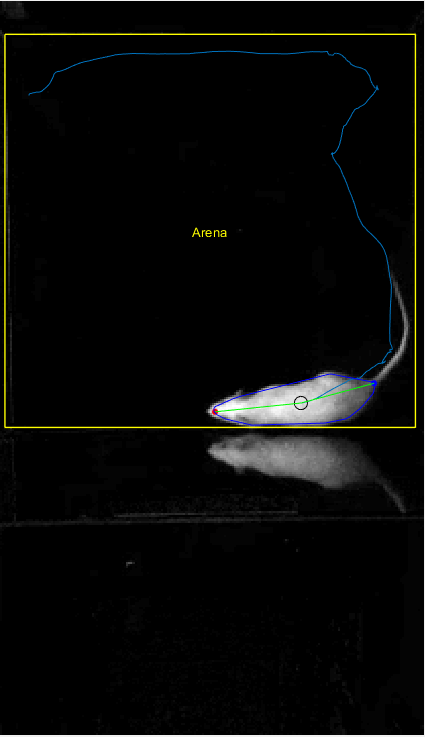

a

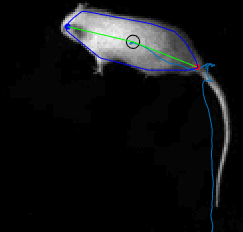

c

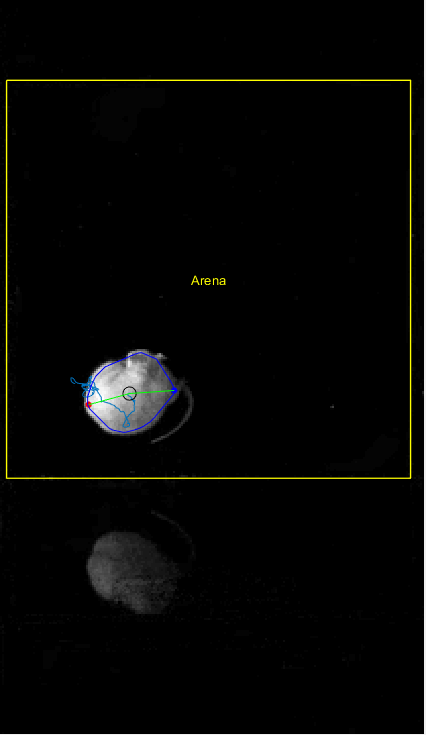

b

**Fig S3. Extremities identification errors.** Examples of tracking when erosion is used to exclude animal’s appendices (tail, legs). (a) The closed dark blue line is the main cluster convex hull after erosion, the open light blue curve marks the centroid (green cross) positions in previous frames and the head (red point) and tail-base (blue point) assignment is successful. (b) A case where the tail is partially visible, and the head/tail assignment is wrong. (c) An example where the head/tail assignment is wrongly swapped.

*TrAQ workflow and GUIs*

To keep TrAQ simple, intuitive, and user-friendly we designed GUIs (Graphical User Interface) for the Tracker and Analyzer modules, requiring minimal intervention to set-up a study and review the data output. The current GUI layouts are shown in Fig S4, reporting the Project Set-Up (Fig S4a) and the Arena Definition (Fig S4b) windows of the Tracker module, the Results Viewer (Fig S4c) and the Region of Interest Definition (Fig S4d) windows of the Analyzer module.

The Project Set-Up window (Fig. S4a) allows the user to modify the tracking settings: first and last video frame to track, threshold and erosion levels. The Arena Definition window (Fig. S4b) allows to select the 2D arena with its physical dimensions. Moreover, for rectangular arenas with sides parallel to the camera FOV, TrAQ implements an automatic rectangular arena recognition algorithm which can be used when the arena borders present a detectable gradient intensity in the background image. The default value for the parameters defined in the Project Set-Up window are easily accessible in the UserSettings.txt file within the software main folder. Editing this file will automatically change all the software default values appearing in the GUIs, reducing the operator intervention when similar videos are to be analyzed.

The Results Viewer window, Fig S4c, allows to review the tracking results. In the same window the user can: (i) set the centroid velocity threshold used to calculate the animal’s activity; (ii) define multiple ROIs within the arena to analyze the animal’s interaction with an object/stimulus (Fig S4d); (iii) activate the head/tail automatic positions correction, defining the maximal admissible displacement value in one frame-step; and (iv) manually correct the head/tail positions in specific frames, an operation that will affect all the subsequent frames since the continuity criterion is adopted from the current frame up to the last one.

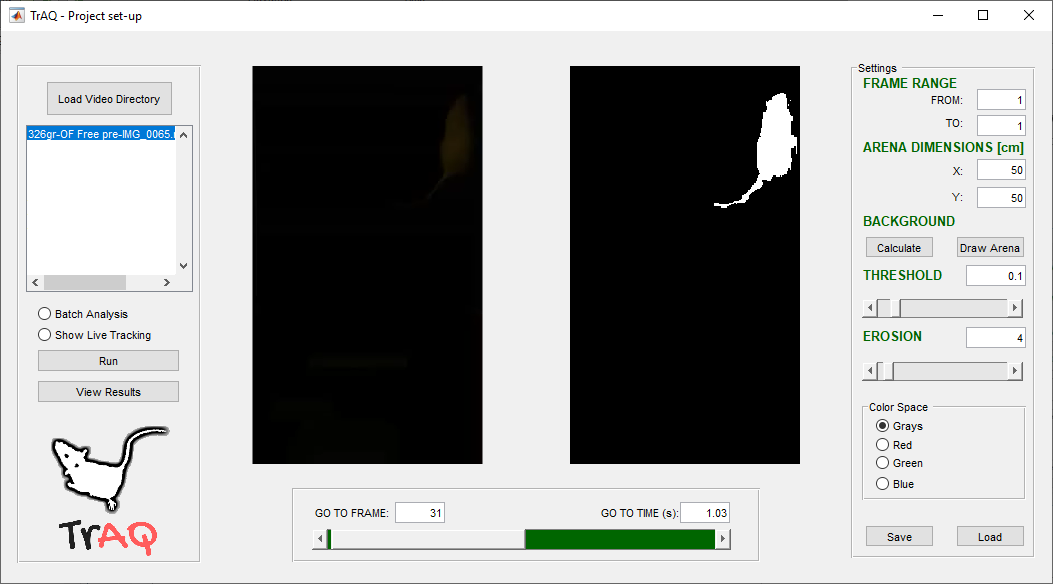

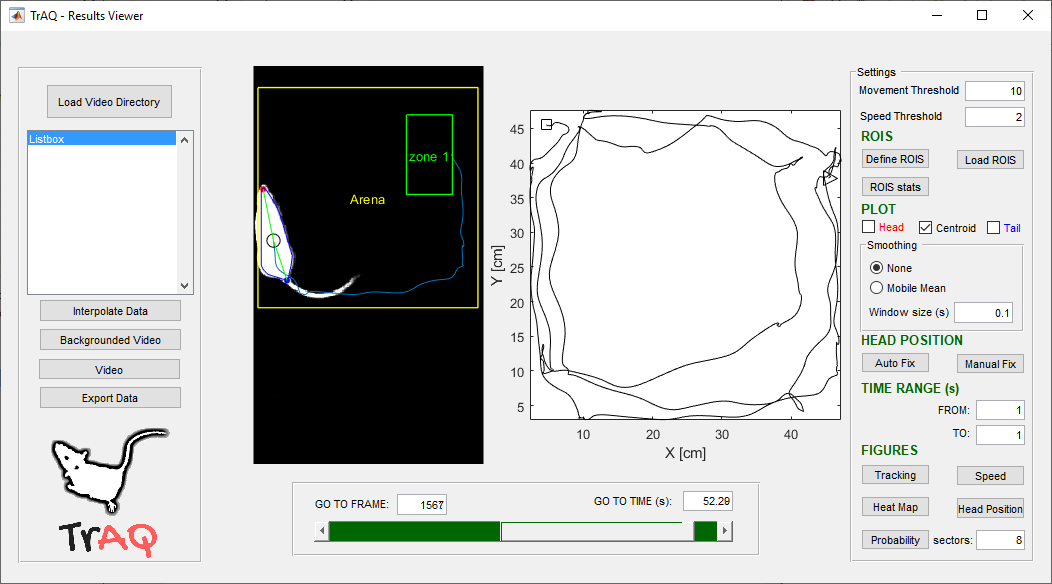

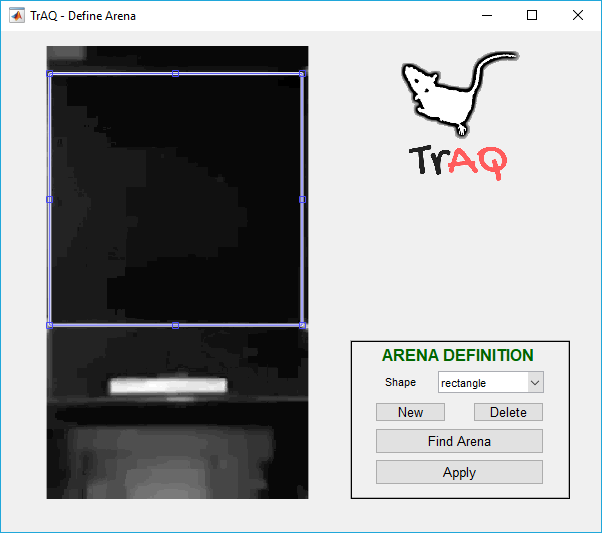

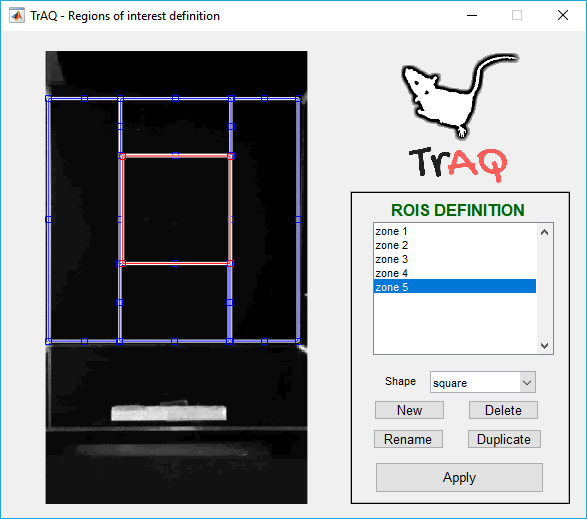

a

b

c

d

**Fig S4. TrAQ GUIs.** TrAQ Graphical User Interfaces (GUIs) windows. Tracker module: (a) Project Set-Up; and (b) Arena Definition. Analyzer module: (c) Results Viewer and (d) Regions of Interest Definition.

The user can customize some parameters or extend the software compatibility by editing the “UserSettings.txt*”* and “video_files.m” files, located in the main software directory.

The UserSettings.txt file contains all the default values of the Project Set-up window operative parameters. Editing the numerical values in the file will change the default values in the software, avoiding the user to modify them every time a new project is started. The available analysis parameters are as follows (the numerical values reported here were used for the 6-OHDA results presented in the manuscript):

arena_x(cm) = 50

arena_y(cm) = 50

Threshold(0-1) = 0.01

Erosion(pixels) = 4

Theta(degrees) = 240

N_frames = 10

Arena x and y are the horizontal and vertical arena dimensions in cm, Threshold is used to generate the binary image, erosion is the neighborhood size (in pixels) used to erode the image (e.g. to remove the tail).

Theta and N_frames are used for rotation counting correction and are defined as the angle (in degrees) and the time window (in frames) used to calculate the rotational velocity threshold.

To extend the software compatibility to new video file format the user can edit the video_files.m file adding a new variable with the needed extension. For example, compatibility to a generic video file extension (.ext) can be added adding the following line:

extfiles = dir([handles.video_dir_text.String filesep '*.ext']);
